# HINGE: Long-Read Assembly Achieves Optimal Repeat Resolution

**DOI:** 10.1101/062117

**Authors:** Govinda M. Kamath, Ilan Shomorony, Fei Xia, Thomas A. Courtade, David N. Tse

## Abstract

Long-read sequencing technologies have the potential to produce gold-standard de novo genome assemblies, but fully exploiting error-prone reads to resolve repeats remains a challenge. Aggressive approaches to repeat resolution often produce mis-assemblies, and conservative approaches lead to unnecessary fragmentation. We present HINGE, an assembler that seeks to achieve optimal repeat resolution by distinguishing repeats that can be resolved given the data from those that cannot. This is accomplished by adding "hinges" to reads for constructing an overlap graph where only unresolvable repeats are merged. As a result, HINGE combines the error resilience of overlap-based assemblers with repeat-resolution capabilities of de Bruijn graph assemblers. HINGE was evaluated on the long-read bacterial datasets from the NCTC project. HINGE produces more finished assemblies than Miniasm and the manual pipeline of NCTC based on the HGAP assembler and Circlator. HINGE also allows us to identify 40 datasets where unresolvable repeats prevent the reliable construction of a unique finished assembly. In these cases, HINGE outputs a visually interpretable assembly graph that encodes all possible finished assemblies consistent with the reads, while other approaches such as the NCTC pipeline and FALCON either fragment the assembly or resolve the ambiguity arbitrarily.

## INTRODUCTION

While genome assembly has been a central task in computational biology for decades, only with the recent advent of long-read technologies has the goal of obtaining near-finished assemblies in an automated fashion become within reach. However, extracting the information present in long error-prone reads in order to reliably resolve repeats is still a challenge (Myers 2016a). Attempts to resolve repeats that are fundamentally unresolvable from the reads at hand - a practice that can be driven by the prospect of a higher N50 score - can lead to incorrect assemblies and ultimately impact downstream scientific analyses. On the other hand, a conservative approach that breaks the assembly at points of seeming ambiguity may fail to produce the longest contigs that can be constructed given the data.

In this sense, an optimal assembler should be one capable of identifying and resolving all, an only those, repeat patterns that are resolvable given the available read data. Equivalently, this objective can be viewed as the construction of an *assembly graph* with the maximum level of repeat resolution that is possible given the data. If a finished assembly of the genome is possible, such a graph would consist of a single cycle (in the case of a single circular chromosome). Otherwise, the next-best objective would be the construction of a *repeat graph* (Pevzner and Tang 2001; Mulyukov and Pevzner 2002) where long repeats are collapsed into a single path. Such paths capture inherent ambiguities about the target genome that cannot be resolved given the data. Thus, constructing the *maximally resolved assembly* graph corresponds to minimizing the number of repeat-induced collapsed segments.

As a prerequisite to this task, one must first understand which repeat patterns can be reliably resolved given the set of reads. Early studies of this fundamental problem appeared in the context of sequencing by hybridization (Ukkonen 1992; Pevzner 1995), and were later extended to shotgun sequencing through the notion of bridging (Bresler et al. 2013). A repeat is said to be *bridged* if at least one read completely contains one of its copies (throughout the paper, we use the word copies to refer to the distinct occurrences of a repeat element). The notion of bridging allows us to define a maximally resolved assembly graph as the graph where only segments corresponding to *unbridged* repeats are collapsed, as discussed in Supplemental Figure S1. The *de novo* construction of such a graph yields the longest contigs that can be reliably constructed, and also describes the plausible arrangements of these contigs in the target genome.

Assembly graphs have been a key component in assembly pipelines since the early days of sequencing projects (Myers et al. 2000). Approaches to assembly graph construction are customarily divided into two categories: de Bruijn graph-based approaches, and overlap-layout-consensus (OLC) approaches. In the de Bruijn framework (Mulyukov and Pevzner 2002; Pevzner and Tang 2001), the set of all *k*-mers is extracted from the reads, and used to build a graph where two *k*-mers that appear consecutively in a read are connected by an edge. This construction has the desirable property that the resulting graph is essentially Eulerian, and repeats longer than *k* base pairs are naturally collapsed into a single path. Furthermore, the graph construction is typically followed by repeat resolution steps using reads that bridge repeats. This allows several de Bruijn graph-based assemblers to produce a maximally resolved assembly graph where only unbridged repeats remain collapsed (Butler et al. 2008; Mulyukov and Pevzner 2002; Peng et al. 2010; Pevzner and Tang 2001).

In the context of third-generation long-read sequencing, however, standard de Bruijn graph approaches have not been as successful as they were in the case of short-read sequencing. Due to the high error rates associated with third-generation platforms, a large number of spurious *k*-mers is created, disrupting the structure of the de Bruijn graph. Recently, the concept of solid *k*-mers was proposed as a way to construct an "approximate" de Bruijn graph on a restricted set of reliable *k*-mers (Lin et al. 2016). However, since overlapping reads only share a handful of solid *k*-mers, the resulting graph lacks the attractive features of de Bruijn graphs. In particular, the Eulerian structure is compromised and repeats are no longer properly collapsed into single paths. Overlap-based approaches, on the other hand, are more robust to read errors since they directly connect reads based on overlaps instead of first breaking them into *k*-mers. In fact, most available long-read assemblers (Chin et al. 2013, 2016; Berlin et al. 2015; Li 2016) are based on the so-called overlap-layout-consensus (OLC) pipeline.

While de Bruijn graphs are Eulerian, overlap graphs are Hamiltonian; i.e., the underlying genome sequence corresponds to a cycle that traverses every node (read) in the graph. In addition to well-known computational challenges (Nagarajan and Pop 2009), the Hamiltonian paradigm does not yield a natural representation of repeat patterns, and the graph is typically riddled with unnecessary edges. In order to combat these issues, the string graph approach (Myers 1995, 2005) was proposed, originally for the Celera assembler (Myers 2016a; Myers et al. 2000), and later adopted by several assembly pipelines (Berlin et al. 2015; Chin et al. 2013, 2016; Li 2016). Built via a *transitive reduction* procedure, the string graph is an overlap graph where the unique, non-repetitive parts of the genome correspond to simple, unbranched, paths. However, long repeats -- both bridged and unbridged -- may result in undesirable graph motifs. In practice, only heuristics are used to combat these motifs, and building a maximally resolved overlap graph is challenging.

## RESULTS

We propose HINGE as a way to build an assembly graph where only the segments corresponding to unbridged repeats are collapsed. This objective, which we refer to as maximally resolved assembly graph, is illustrated in Figure 1(a)-(e). As depicted in Figure 1(f), this goal is naturally achieved in a de Bruijn graph framework, but not within an overlap graph-based framework due to the motifs created by long repeats. HINGE seeks to simultaneously attain the error resilience of overlap graph-based approaches and the appealing graph structure and optimal repeat resolution capability of de Bruijn graphs. Next we briefly outline the main algorithmic innovations that allow HINGE to achieve this goal, and present results on several datasets.

**Figure 1:**
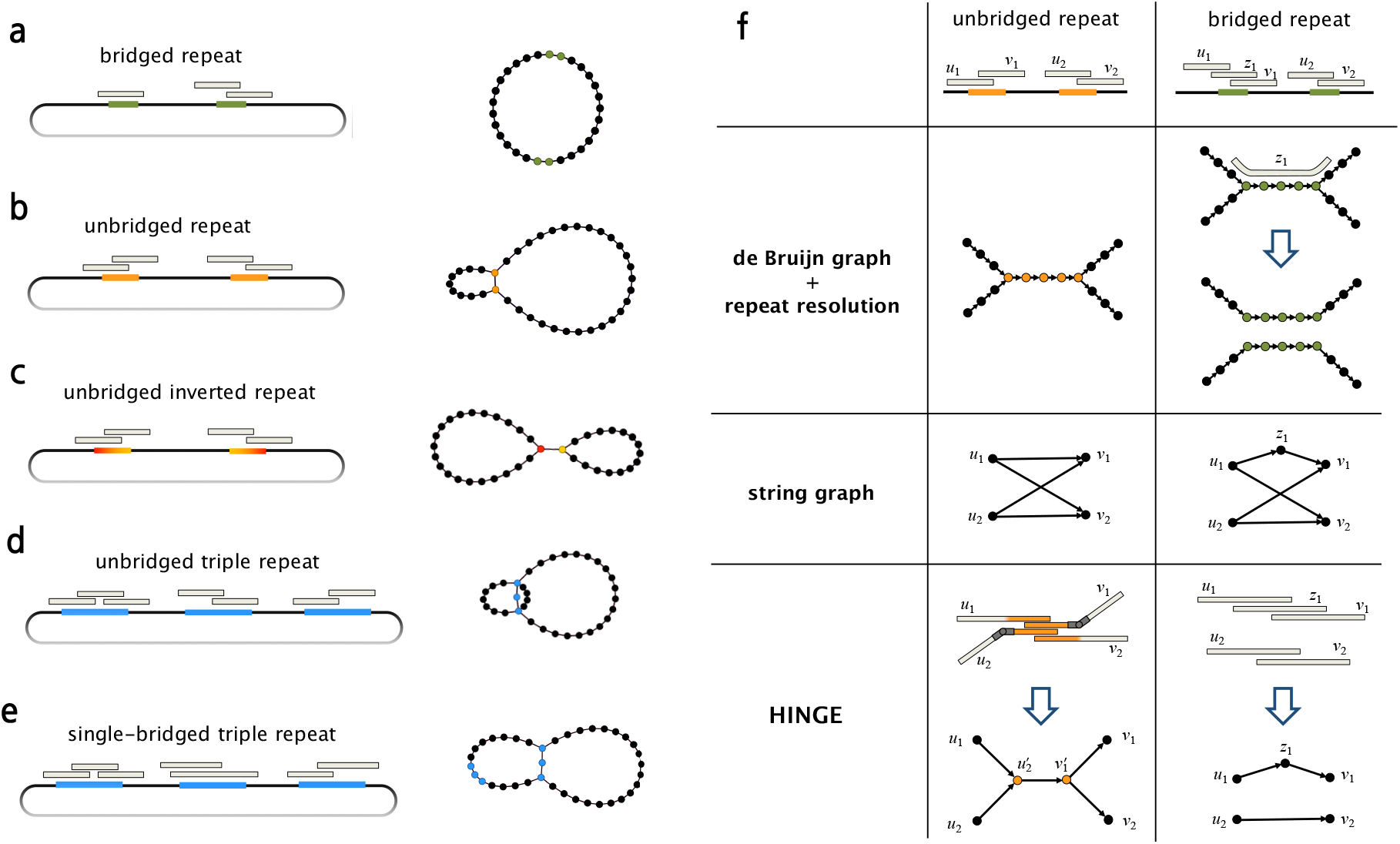
The goal of HINGE is to produce a maximally resolved assembly graph, where repeats that are bridged by the reads are not collapsed, and repeats that are unbridged are collapsed in a natural way, similar to what is achieved with de Bruijn graphs, **(a)** If at least one of the two copies of a repeat is bridged (green segments), the maximally resolved assembly graph should separate the two copies. In **(b-e)**, we illustrate an unbridged repeat, an unbridged inverted (i.e., reverse-complemented) repeat, an unbridged triple repeat, and a single-bridged triple repeat, and the assembly graph obtained by collapsing segments corresponding to unbridged repeats. Notice that in **(b,e)** the graph admits a single traversal and can be further resolved, while in **(c,d)** the graph admits two distinct traversals and cannot be further resolved (see Supplemental Figure S15). **(f)** The representation of a bridged and an unbridged repeat in the de Bruijn graph approach, in the standard string graph approach, and according to HINGE. The de Bruijn graph approach collapses the repeated segment, which allows a natural repeat resolution step if a bridging read is found. The representation in the string graph (if there is no read entirely contained in the repeat) is an hourglass-like motif. HINGE emulates the de Bruijn graph layout, but in an overlap graph framework.

## ALGORITHMIC CONTRIBUTIONS

HINGE is an assembler that follows the Overlap-Layout-Consensus (OLC) paradigm. Its main algorithmic innovation lies in how it exploits the alignments obtained in the Overlap phase in order to identify resolvable repeats and construct the graph layout in a repeat-aware fashion. Next we describe the main ideas that go into the Layout step. We defer a description of the overlap and consensus steps to the METHODS section.

### Repeat annotation and hinging reads

HINGE utilizes the alignment information obtained in the Overlap step in order to equip some of the reads with *hinges.* Hinges are placed at the beginning and end of unbridged repeats, and will ultimately lead to bifurcations on the graph, as illustrated in Figure 1(f). The first step towards hinging the reads, as illustrated in Figure 4(a), is to find sharp gradients in the number of alignments on a read and annotate them as beginning or end of repeats. Next, we identify reads that bridge a repeat by finding reads that have both an annotation for the beginning of a repeat and an annotation for the end of the same repeat.

Finally, we spread the information of which repeats are bridged to other reads through a procedure that we term the *Contagion algorithm* (see Methods and Figure 5).

### Hinge-aided greedy overlap graph construction

The Contagion algorithm allows HINGE to place exactly one in-hinge and one out-hinge on the reads that originated from unbridged occurrences of a repeat. HINGE can then create a sparse overlap graph by using a hinge-aided greedy graph construction. In essence, we pick a best predecessor and a best successor for each read, as in the classical greedy algorithm (Tarhio and Ukkonen 1988) or in the best-overlap-graph approach (Miller et al. 2008a). However, since our reads are hinged, we also allow a read’s successor or predecessor to be the *interior* of another read, as long as the match starts on a hinge. When this occurs, a bifurcation is formed on the graph, corresponding to the beginning or the end of an unbridged repeat.

As illustrated in Figure 1(f), this hinge-aided approach allows us to obtain the attractive properties of a de Bruijn graph layout, but within the OLC framework. A comparison with the traditional greedy approach is provided in Supplemental Figure S2. We point out that for higher fold repeats, where a subset of the copies may be unbridged, a more careful handling of hinges is required, and that is achieved using a new procedure that we call *Poisoning*, described in the METHODS section and in Figure 6.

## VALIDATION OF HINGE ON DATASETS WITH GROUND TRUTH

In Supplemental Figures S3, S4, S5, and S6, we present validation results on simulated datasets. We created sequences with specific patterns of repeats, and simulated long error-prone reads, using the DAZZ-DB simulator. We then verified that, when run on these datasets, HINGE produces a maximally resolved assembly graph. In Supplemental Figure S7, we validate the structural integrity of our assembly on an Oxford Nanopore R9 *E. coli* dataset. In Supplemental Figure S8, we validate the structural and sequence integrity of our assembly on a PacBio *Saccharomyces cerevisiae* dataset.

In Supplemental Table S2, we present validation results on *E. coli* datasets produced by PacBio and Oxford Nanopore sequencers. In both of these cases, HINGE produces a single circular contig and there is no misassembly. We also compare our assembly with the assembly produced by the NCTC pipeline (HGAP followed by Circlator) on 10 randomly selected datasets. We verify that the assemblies agree and have high identity scores in all cases.

## EVALUATION ON THE NCTC DATABASE

We evaluated HINGE on the 997 bacterial genomes of the NCTC 3000 database that were publicly available at the time of writing this manuscript (Wellcome Trust Sanger Institute 2016). The accession number for these datasets is provided in Supplemental Tables S1 and S3. Each of these datasets consists of PacBio SMRT long reads with coverage depths mainly in the range 30x to 80x. While the repeat complexity is relatively mild in bacterial genomes, we chose to evaluate HINGE on these datasets for two reasons: it allows us to carefully verify whether the HINGE assembly graphs satisfy our goal of maximal repeat resolution, and it allows us run experiments on a large number of datasets, thus avoiding overfitting.

The current NCTC manual assembly pipeline uses the HGAP assembler (Chin et al. 2013) to produce a list of contigs, and Circlator (Hunt et al. 2015) to circularize contigs. The assembly graphs produced by HINGE with no parameter tuning for each of these datasets are available online (Kamath et al. 2016) and in Supplemental Table S4, along with the contig statistics of the NCTC pipeline results, and the assembly graph produced by Miniasm (Li 2016). We point out that other state-of-the-art assemblers, in particular FALCON (Chin et al. 2016), have runtime above one order of magnitude greater than HINGE (see Supplemental Figure S1 1), making a comparison on the entire NCTC database computationally prohibitive.

For 822 of the 997 available datasets, HINGE produced a finished non-fragmented assembly graph, with additional isolated small plasmids in many cases. In 40 of these datasets, HINGE identifies unresolvable repeats, and the final graph admits distinct traversals (See Table 1). In order to compare our results with those obtained by the NCTC manual pipeline, we restricted our attention to those datasets for which NCTC reports the results of their assembly. As shown in Table 2, even without a circularization tool, HINGE obtains significantly more finished assemblies than the NCTC pipeline.

**Table 1:** Finished assemblies on all available NCTC datasets and comparison with Miniasm. Given the output graph of HINGE we classify the assembly into four categories. A *finished circular assembly* corresponds to a case where all nodes (small plasmids excepted) lie on a single circle. A *finished circular assembly with multiple traversals* corresponds to a graph where all nodes can be visited by a circular path, but there is more than one such path. We point out that we classify such an output as finished because such a graph can be seen as simultaneously capturing a few (usually two) assemblies, all of which would be considered finished according to the previous rule. A finished assembly is said to *lack circularization* if a single non-circular path can traverse all nodes on the graph (small plasmids excepted). If the graph produced by HINGE does not fall into the previous three categories, we classify it as a mis-assembly/fragmented assembly. As reliable hinge placement requires a reasonable coverage depth, we also considered restricting our attention to the datasets with average coverage depth above 40. We note that Miniasm needs a circularization tool to circularize assemblies, and hence we report a Miniasm assembly as finished if it has only one contig longer than 200 kbp and fewer than ten contigs shorter than 200 kbp. The graph produced by HINGE and Miniasm for all these cases can be found at http://web.stanford.edu/~gkamath/NCTC/report.html and in Supplemental Table S4 along with the corresponding classification. As can be seen on this report, the rule for determining when a Miniasm assembly is finished is often quite lenient.

| | Coverage $\geq 40\times$ | | | All coverages | | |
| --- | --- | --- | --- | --- | --- | --- |
| Number of NCTC datasets | 816 |  |  | 997 |  |  |
| **HINGE** finished circular assembly (single traversal) | 631 | 691 | 729 | 690 | 782 | 822 |
| **HINGE** finished assembly (lacking circularization) | 60 |  |  | 92 |  |  |
| **HINGE** finished circular assembly (multiple traversals) | 38 |  |  | 40 |  |  |

**Table 2:** Finished assemblies on NCTC datasets where NCTC manual pipeline results are reported. In this table we restrict the datasets considered in Table 1 to only those for which NCTC reports a result for comparison. The finished assemblies for the NCTC manual pipeline correspond to the cases where they report one chromosomal contig or two chromosomal contigs (since species such as *Vibrio fluvialis* and *Ochrobactrum anthropi* are known to have two chromosomes). We point out that while a circularization tool (Circlator) is used in the NCTC pipeline, we do not have a circularization finishing step and only report the output of HINGE using default configurations.

| | Coverage $\geq 40\times$ | | All coverages | |
| --- | --- | --- | --- | --- |
| Number of NCTC datasets | 688 |  | 834 |  |
| NCTC manual pipeline finished assemblies | 517 |  | 592 |  |
| **Miniasm** finished assembly (not circularized) | 513 |  | 592 |  |
| **HINGE** finished circular assembly (single traversal) | 531 | 583 | 583 | 660 |
| **HINGE** finished assembly (lacking circularization) | 52 |  | 77 |  |
| **HINGE** finished circular assembly (multiple traversals) | 33 |  | 33 |  |

## ANALYSIS OF HINGE ASSEMBLY GRAPHS

Among the cases where HINGE produces an assembly graph with multiple traversals, we find many examples where the intuitive layout of the graph produced by HINGE resembles the idealized cases in Figure 1(a-e), and allows one to visually assess the unresolvable repeat pattern in the genome. Next, we analyze three such cases in depth, and compare the graph produced by HINGE with the contigs produced by the NCTC pipeline. We see that by focusing on obtaining a maximally resolved assembly graph rather than large contig N50 values, HINGE prevents several mis-assemblies the NCTC pipeline incurred. In Supplemental Figure S9, we present nine additional such cases. In Supplemental Figure S1 0, we present several cases where HINGE resolves all repeats, producing a finished circular assembly, while the NCTC pipeline instead fragments the assembly. In addition, in Supplemental Figures S12, S13, and S14 we provide the same comparisons but with FALCON (Chin et al. 2016) instead of the manual NCTC pipeline.

In Figure 2(a), we examine NCTC11022 *(Escherichia coli).* In this example, the incorrect resolution of a 20 kbp unbridged repeat by the NCTC pipeline (see Supplemental Figure S1 6) causes the circular chromosomal contig to lose a 780 kb segment, returned as a separate contig. By first collapsing this repeat and then resolving it due to the existence of a unique traversal of the graph, HINGE produces a single large chromosomal contig of length 5.1 Mbp. The nodes in the HINGE graph are colored according to the position the corresponding reads align to in the NCTC pipeline contigs.

**Figure 2:**
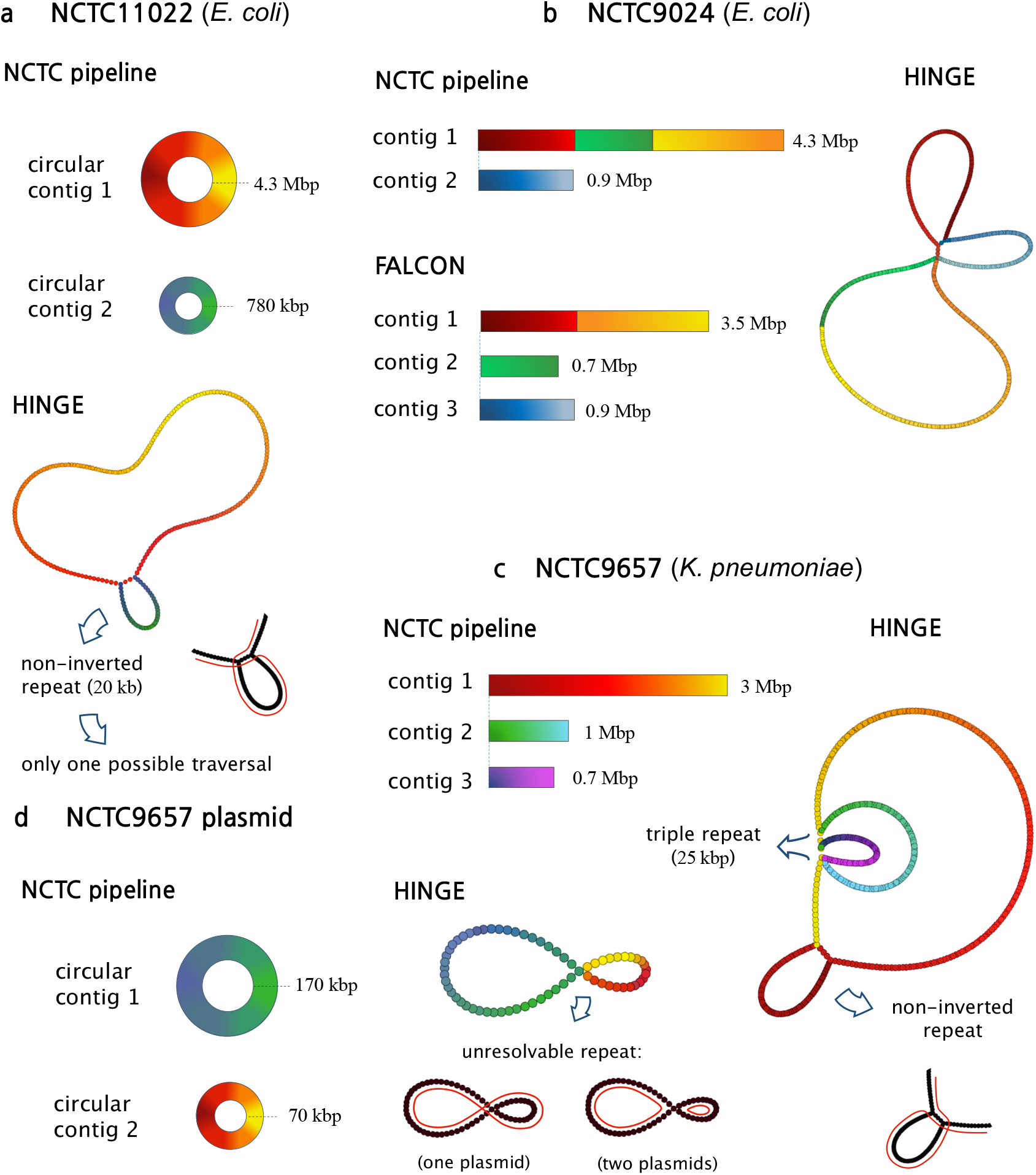
Analysis of HINGE graphs on selected datasets. By identifying unbridged repeats, collapsing them, and then performing resolutions based on uniquely traversable loops, HINGE prevents mis-assemblies and produces a user-friendly interpretable assembly graph. We color the graph nodes according to their corresponding position on the NCTC pipeline contigs. (a) On NCTC11022, HINGE identifies an unbridged repeat, which is later resolved, (b) On NCTC9024, HINGE identifies an unbridged triple repeat (with one inverted copy), which cannot be resolved due to the existence of three distinct traversals of the graph, (c) HINGE identifies an unbridged triple repeat, (d) HINGE identifies an unresolvable repeat shared by two small plasmids.

On the NCTC9024 dataset *(Escherichia coli)* (Figure 2(b)), the NCTC pipeline returned two long contigs, one of 4.3 Mbp and one of 0.9 Mbp. The HINGE graph emphasizes the existence of a triple repeat which, upon further inspection (See Supplemental Figure S1 7), is seen to be of length 20 kbp, unbridged, and with one inverted copy. Even though this repeat is unbridged, both the NCTC pipeline and FALCON resolve one of its copies, but in distinct ways. As we point out in Supplemental Figure S9, incorrect resolution of an inverted repeat can produce a false inversion of a long contig. In fact, the NCTC assembly and the FALCON assembly disagree on the orientation of the yellow-to-orange segment, and one of them must be creating an incorrect inversion of more than 1 Mbp (the orange-to-yellow segment). By collapsing the repeat, HINGE avoids a potential mis-assembly.

In Figure 2(c), we consider NCTC9657 *(Klebsiella pneumoniae).* In this example, the NCTC pipeline returned seven unidentified contigs (three large ones), but HINGE returns a single large chromosomal connected component, and three small plasmids. In this case, HINGE produces a graph motif characteristic of an unbridged triple repeat, similar to Figure 1 (d). As shown by a coverage analysis in Supplemental Figure S1 8(a), this is indeed a triple repeat and contig 1 of the NCTC pipeline incorrectly resolves it, creating a mis-assembly. In addition, we examine the plasmids produced by the NCTC pipeline in Figure 2(d), and note that two of them share an unbridged repeat (see also Supplemental Figure S1 8(b)). Therefore, there are two possible resolutions (two plasmids or a single, longer, plasmid), and HINGE keeps them merged on the graph to retain this unresolvable ambiguity. In Supplemental Figure S1 2, we verify that the performance of FALCON (Chin et al. 2016) on the examples in Figure 2 (a) and 2(c) is similar to that of the NCTC pipeline.

As illustrated by these examples, HINGE seeks to construct a user-friendly, informative, overlap graph as its main output, as opposed to most OLC assemblers, which employ assembly graphs in their inner workings (Berlin et al. 2015; Chin et al. 2013, 2016) but focus on outputting a list of contigs. To the best of our knowledge, Miniasm (Li 2016) is the only other assembler to produce a graph as the main assembly output. However, Miniasm is based on the string graph paradigm, which does not achieve the graph layout HINGE strives for as we empirically observe (Kamath et al. 2016).

## DISCUSSION

With HINGE, we introduce a new approach to constructing assembly graphs in a repeat-aware fashion. While other state-of-the-art assemblers do attempt to identify bridging reads (sometimes referred to as spanning reads) and resolve the corresponding repeats, this is usually done as a post-processing step on the graph. HINGE, on the other hand, seeks to identify repeats and determine whether they should be collapsed on the graph prior to the actual construction. This way, HINGE avoids having to identify and correct graph motifs (such as the ones created by the string graph as shown in Figure 1(f)) in a post-processing phase, which can be difficult due to spurious and missing edges caused by the high error rates of long-read sequencing technologies and by chimeric reads.

In order to reliably achieve this repeat-aware graph layout, several new conceptual ideas were introduced in HINGE. First, a repeat annotation step is responsible for identifying the beginning and end of repeats and which reads bridge some repeat. However, this type of local information is not sufficient for the construction of a maximally resolved assembly graph. Therefore, this information must be spread to other reads, which is accomplished with our *Contagion* algorithm.

Once the bridging information is known globally, HINGE utilizes a hinge-aided greedy construction of the graph. This is also different from most state-of-the-art long-read assemblers, which rely on the string graph paradigm. Our approach bears similarities with the Best Overlap Graph approach in its goal of constructing a sparse overlap graph, but takes advantage of hinges as a way to achieve this goal with maximal repeat resolution. Finally, the sparse nature of the constructed graphs allows HINGE to identify loops that admit a single traversal and can thus be resolved. The conceptual contributions of HINGE are discussed in more detail in the METHODS section.

As an OLC assembler, in order to produce high quality assemblies, HINGE relies on good Overlapping and Consensus modules. In its current implementation, HINGE was designed to work with the output of DALIGNER (Myers 2014), and the consensus is performed using a variant of the consensus module of FALCON (Chin et al. 2016) together with a straightforward majority-vote finishing step. These choices are not essential to the workings of our pipeline. Therefore, integrating HINGE with other overlapping tools such as MHAP or Minimap can be done if different levels of alignment sensitivity or memory usage are required. Similarly, different consensus and polishing modules such as Quiver (Chin et al. 2013) and Racon (Vaser et al. 2016) can be used, according to the desired point in the accuracy-computation tradeoff.

Through a novel approach to repeat resolution and graph representation, HINGE brings a fresh perspective to the assembly problem. By focusing on the construction of a maximally resolved assembly graph in a user-friendly fashion, HINGE is well aligned with the recent push for the standardization of graph references, as opposed to the traditional contig representation. The HINGE graph is a natural representation of a set of possible assemblies, and is amenable to further repeat resolution, which can be attempted using additional long-range information such as paired-end reads, Hi-C reads, or by leveraging biological insight. Finally, we point out that while the repeat complexity is relatively mild in the bacterial genomes we consider (as evidenced by the large number of finished assemblies), it is much more severe in higher organisms (Koren et al. 2013). This highlights the importance of the careful treatment to repeats carried out by HINGE and the value of the proposed method to genome assembly.

One important aspect regarding the notion of maximal repeat resolution is that it assumes that long contiguous matches identified in the read alignment step must correspond either to the same segment on the genome or to repeats whose copies are similar enough that they should be merged in the graph. However, there may still be a small level of divergence between these copies that is below the sequencing error rates and cannot be detected by the aligner. In principle, this divergence may allow a final “phasing” or “unzipping” step, similar to what is used in FALCON-Unzip (Chin et al. 2016a), to resolve these repeats. Utilizing these small levels of divergence to phase or to score the different traversals of a repeat according to their likelihood is a future direction for improvement of the HINGE pipeline.

## METHODS

The HINGE assembly pipeline is an OLC pipeline designed to assemble long reads. The overall workflow is depicted in Figure 3, and is explained in detail in this section. As the default parameters and auxiliary tools were selected to optimize the pipeline for PacBio reads, we focus the discussion on this setting.

**Figure 3:**
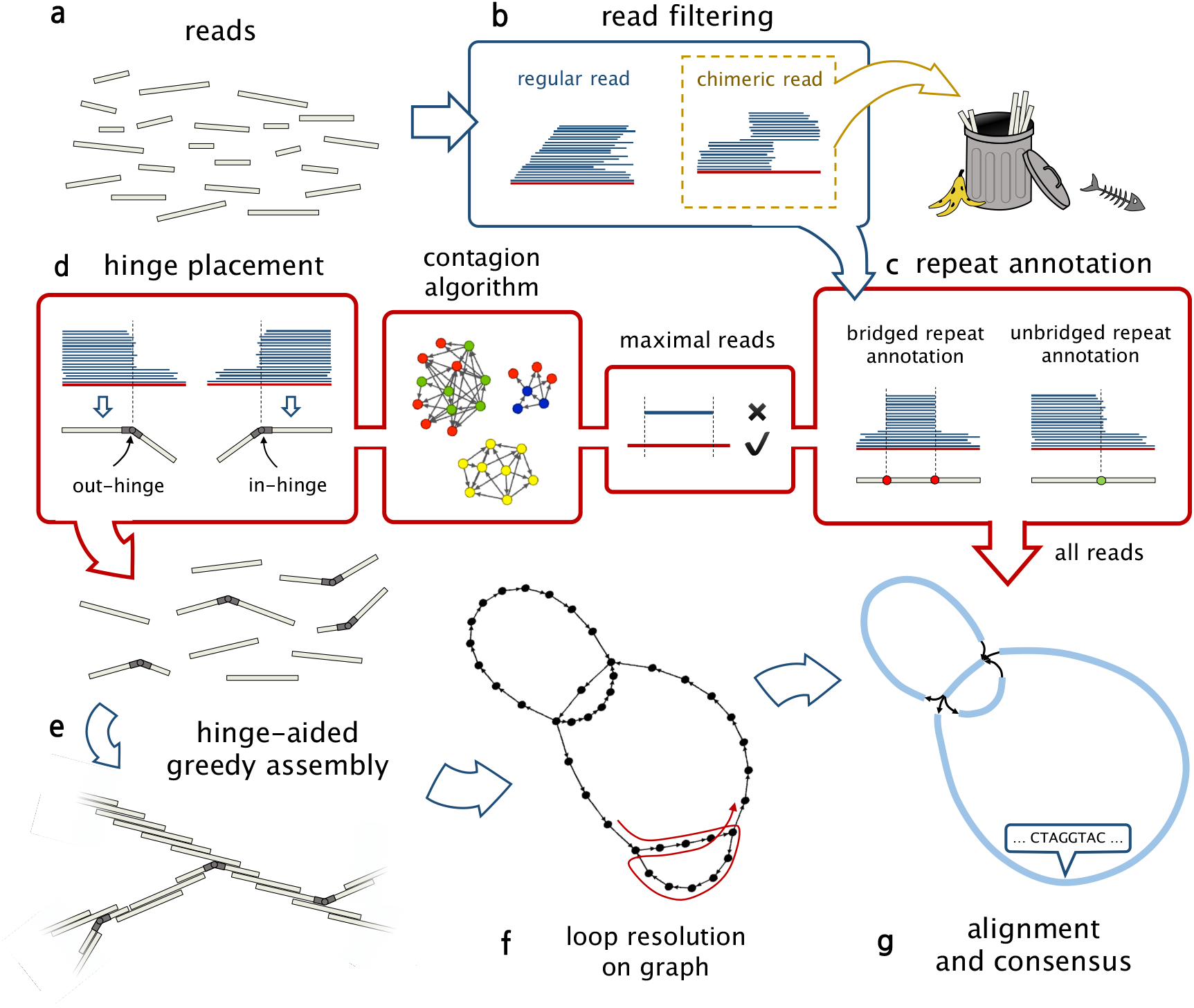
HINGE pipeline: **(a)** The input to the HINGE pipeline is a set of long error-prone reads, **(b)** Chimeric reads are detected through their pile-o-grams, and are discarded, **(c)** The beginning/end of repeats are annotated on the reads. This is done by detecting a sharp increase/decrease in the number of alignments on a read. The repeat annotations are also identified as bridged or unbridged, **(d)** Maximal reads (i.e., reads that are not contained in other reads) are selected, and fed to the contagion algorithm, which is responsible for spreading the information about which repeats are bridged to all the reads, allowing us to place exactly one in-hinge and one out-hinge on the reads that originated from unbridged occurrences of a repeat, **(e)** The set of maximal reads (some of which are no hinged) is the input to the hinge-aided greedy assembly, **(f)** After obtaining the read-overlap graph, we resolve repeats that admit only one traversal, **(g)** Finally, by mapping all the reads onto the resulting overlap graph, we use standard consensus methods to generate contigs.

## READ DATABASE AND ALIGNMENT

We use DAZZ_DB (Myers 2016b) to maintain a database of the PacBio reads. We use DALIGNER (Myers 2014) to obtain pairwise alignments between all reads. We point out that HINGE does not heavily rely on specifics of the DALIGNER output, and can be adapted to work with other aligners as well.

## NO INITIAL ERROR-CORRECTION STEP

Unlike most available long-read assembly pipelines, HINGE bypasses an initial error correction step. To the best of our knowledge, Miniasm (Li 2016) is the only other OLC assembler that dispenses with this step. Abruijn (Lin et al. 2016) also has no error-correction step, though it is not based on the OLC paradigm. The idea of error-correction-free assembly was also utilized in (Lien et al. 2016; Tørresen et al. 2016). The fact that long-read aligners like DALIGNER (Myers 2014) can obtain pairwise alignments at error rates around 15% allows us to use this approach, and defer the error correction to the final consensus step.

## CHIMERIC READ FILTER

Chimeric reads are the result of a sequencing error, and are usually made up of multiple segments that originate from different parts of the genome. If not properly handled, these reads create mis-assemblies, and different techniques have been put forward to detect chimera (Miller et al. 2008a; Li 2016). HINGE's chimera filter unit is the first place in the pipeline where the visualization provided by pile-o-grams (Supplemental Figures S19 and S20) is useful. We mark a read segment as chimeric if the set of reads aligned to it undergoes an abrupt change. On the pile-o-gram, as shown in Figure 4(b), one sees a clear discontinuity in the set of alignments (blue segments) of a read. We also mark a read segment as chimeric if the number of matches goes below a fixed threshold. For each read, we keep the longest segment without any chimeric segments. If this segment is shorter than a threshold, we discard the read completely.

**Figure 4:**
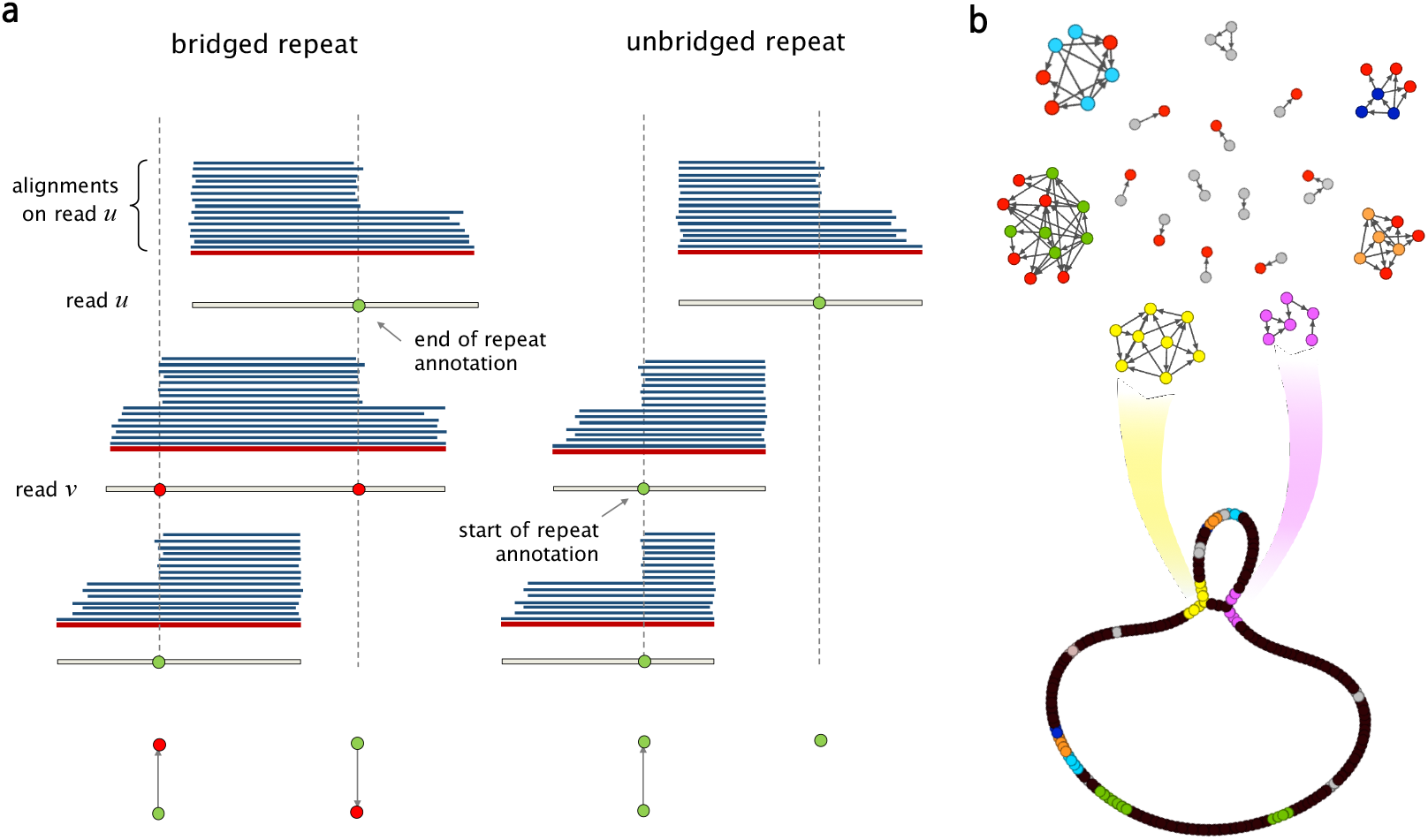
**(a)** Sharp changes in the number of alignments give rise to repeat annotations on each read. If a read is verified to bridge a repeated, as in the case of read v, the corresponding read annotations are marked as such (shown as red nodes), **(b)** The Contagion graph is formed by having all repeat annotations as nodes, and using edges to mark annotations that correspond to the beginning (or end) of the same repeat. As illustrated here for NCTC11022, connected components with no bridged repeat annotations will give rise to hinged reads, which leads to bifurcations on the graph. The repeats corresponding to other connected components stay resolved in the graph.

## REPEAT ANNOTATION

One of the main distinctive features of HINGE is a pre-assembly step responsible for annotating the beginning and the end of repeats on the reads. These repeat annotations will later be used for placing hinges on the reads, which in turn will be instrumental in the graph layout step. The repeat annotation is done by detecting the start/end of a large number of matches on a read. On the pile-o-gram (Supplemental Figures S19 and S20), this visually corresponds to a large pile of matches starting/ending at the same point, as shown in Figure 3(c). We note that relying on coverage gradients rather than coverage itself makes HINGE immune to coverage fluctuations.

We then verify whether the repeat annotation corresponds to a repeat that is bridged by that read. Intuitively, one could attempt to do this by identifying both a sharp increase and a sharp decrease in the number of matches on a read. However, as it turns out, such an approach can fail in the presence of more complex repeat patterns such a repeat within a longer repeat (see Supplemental Figure S1 9(e) for an illustration). Therefore, a more careful processing of the matches on a read is needed to identify the bridging condition. HINGE determines the bridging condition by checking whether most of the matches starting on a repeat annotation also end on a repeat annotation. If that is the case, the repeat is assumed to be bridged, and the annotation is flagged as such (red annotations in Figure 3(c)). Thus at the end of this step we have repeat annotations on all reads, and these annotations are labeled as bridged/unbridged according to the *local* information provided by the reads’ alignments. The next step, the Contagion algorithm, is applied to this set of annotated reads, after we filter out reads that are fully contained in other reads (keeping only maximal reads).

## THE *CONTAGION* ALGORITHM

Notice that this local information about the bridging of repeats may be misleading. For example, the pile-o-gram of read *u* in Figure 4(a) may suggest that *u* lies partially on an unbridged repeat. However, that repeat might still be bridged by a different read, as in the case of read *v* in Figure 4(a). Therefore, HINGE proceeds to “spread” the local bridging information of each read to other reads using the *Contagion* algorithm. At a high level, this algorithm can be thought of as constructing a contagion graph (see Figure 4(b)) with nodes being the repeat annotations, and edges between repeat annotations that correspond to the beginning (or end) of the same repeat (possibly from different copies of the same repeat). Annotations corresponding to the beginning/end of the same repeat are identified based on alignments: if two reads have an annotation corresponding to the beginning (resp. end) of a repeat and have matching segments after (resp. before) the annotation, the two annotations are connected in the graph. The edge points in the direction of the read that extends the most into the repeat. Moreover, repeat annotations that have been identified as the beginning/end of a bridged repeat are marked as such (red nodes in Figure 4(b)).

As illustrated in Figure 4(b), this graph has two connected components for each repeat (the yellow and pink components correspond to the beginning and end of an unbridged repeat). In this graph, repeat annotations corresponding to bridged repeats are thought of as “infected” (shown as red nodes) and can spread the “bridging condition” to the other repeat annotations in the same connected component. If a connected component does not contain any infected repeat annotation, it corresponds to the beginning/end of an unbridged repeat and will eventually lead to a bifurcation on the graph (see Hinge-Aided Greedy Layout), as shown in Figure 4(b).

The Contagion algorithm processes the contagion graph to kill repeat annotations that will not be useful in the overlap graph construction. In particular, this infecting and killing process performs two tasks: (1) repeat annotations corresponding to bridged repeats should cause other annotations corresponding to the same repeat to also be marked as bridged and ultimately killed; (2) if two repeat annotations correspond to the beginning (or end) of the same repeat, the one extending the most into the repeat should be kept, while the other one should be killed. This global processing of the repeat annotations and bridging condition is important so that ultimately we only place one in-hinge and one out-hinge for each unbridged repeat (on the sink node of the corresponding connected component).

In more detail, the contagion algorithm comprises three steps. In the first step, we remove all annotations whose connected component on the contagion graph is small. For instance, the gray-colored nodes in Figure 4(b) correspond to small connected components that are deleted.

Typically, these small components are the result of imprecise placement of repeat annotation on reads, which then lead to them not matching other repeats annotations that correspond to the beginning/end of the same repeat. Hence, deleting these small components prevents us from creating multiple hinges corresponding to the beginning/end of the same repeat.

In the next step, we look for pairs of repeat annotations connected by an edge in the contagion graph (i.e., corresponding to the beginning/end of the same repeat) and such that the corresponding reads have an overlap. (Notice that by an overlap, we mean a match between the suffix of a read and the prefix of another read. If the match instead occurs at the interior of at least one of the reads, we refer to it as an internal match.) For every such pair, we kill the annotation on the read that extends the least into the repeat. For an illustration of this step, consider the two unbridged repeats in Figure 5(a). The reads covering the start of each repeat (*u*_1_, *u*_2_, *u*_3_) have a start-repeat annotation at the start of the repeat. The reads covering the end of each repeat (*v*_1_ *v*_2_, *v*_3_) have an end-repeat annotation at the end of the repeat. As shown in Figure 5(b), the start-repeat annotation on *u_2_* is killed by the start-repeat annotation on *u*_1_ because *u*_2_ and *u*_1_ have and *u*_2_ extends more into the repeat. Similarly, the end-repeat annotation at *v*_2_ is killed due to the overlap with *v*_1_. At the end of this step, we have that exactly one read covering each copy of an unbridged repeat has a start-repeat annotation on it (and exactly one read has an end-repeat annotation on it for each copy of the repeat). In addition, we point out that when a read has its repeat annotation killed by an annotation from a bridged repeat, we mark this annotation as "poisoned". The reason for the term is that a poisoned read would be “deadly” for a standard greedy assembly algorithm, as it would lead to a mis-assembly.

**Figure 5:**
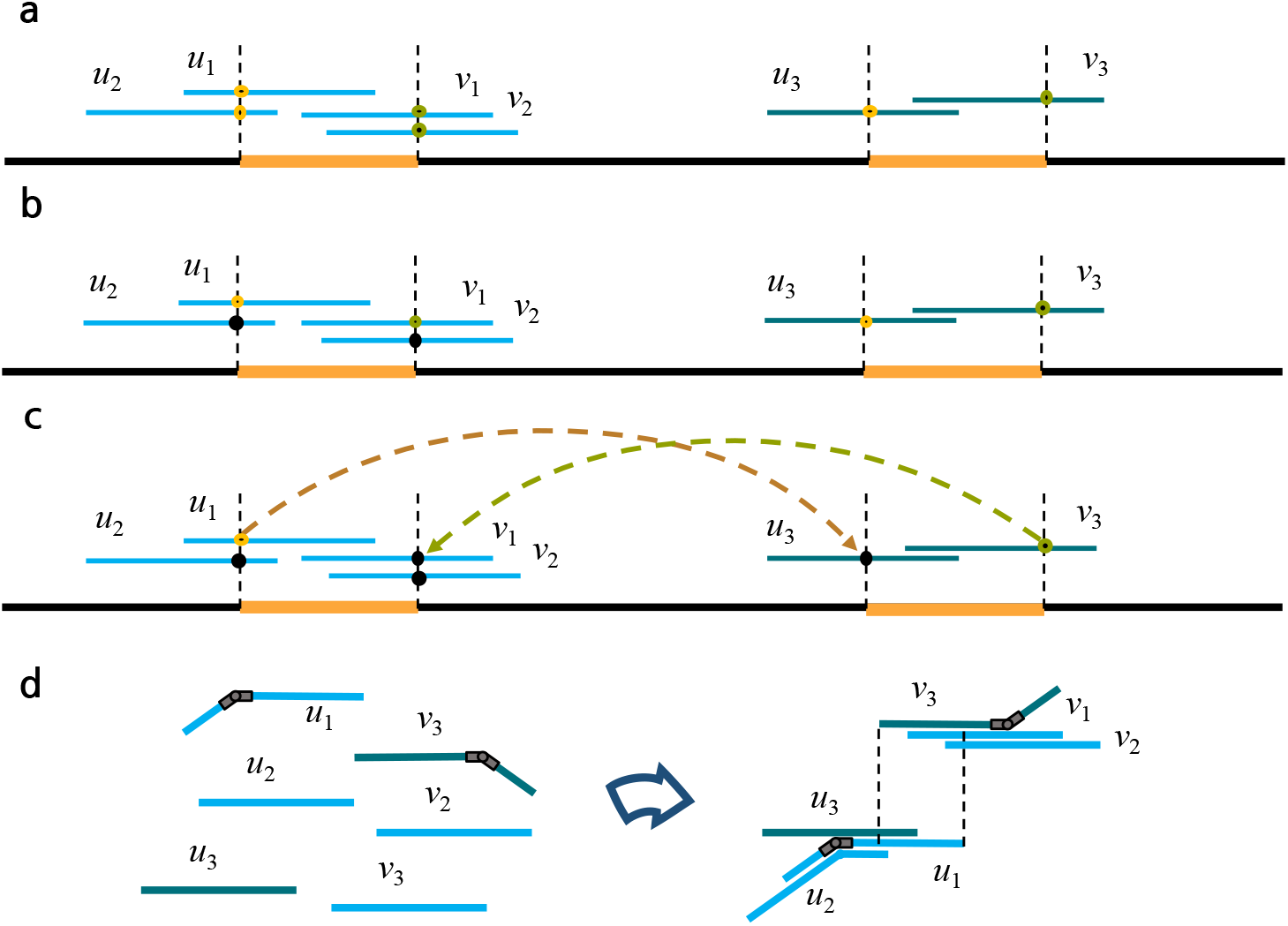
The *Contagion* Algorithm: **(a)** Two unbridged repeats are shown as orange segments, **(b)** The contagion algorithm first kills the start-repeat annotation at *u*_2_ (due to its overlap with *u*_1_) and the end-repeat annotation at *v*_2_ (due to its overlap with *v*_1_). **(c)** The contagion algorithm then kills the start-repeat annotation on *u*_3_ (due to its internal match with *u*_1_), and the end-repeat annotation on *v*_1_ (due to its internal match with *v*_3_). **(d)** Finally, an in-hinge is placed on *u*_1_ and an out-hinge is placed on *v*_3_. During the hinge-aided greedy assembly step, hinges allow *u*_2_ and *u*_3_ to choose the in-hinge at *u*_1_ as their successor. Similarly, *v*_1_ and *v*_2_ pick the match starting at the out-hinge on *v*_3_ as their predecessor match.

The third step of the Contagion algorithm is similar to the second step but instead of looking for matching annotations whose reads have an overlap, we look for matching annotations whose reads have an internal match. For every such pair of annotations, we keep the one on the read that extends the most into the repeat, and kill the other one. As illustrated in Figure 5(c), this causes the start-repeat annotation on *u*_3_ to be killed by the start-repeat annotation on *u*_1_ and the end-repeat annotation on to be killed by the end-repeat annotation on v_3_. At the end of this step, we have one in-hinge on *u*_1_ and one out-hinge on *v*_3_. We point out that in this step we only consider non-poisoned reads.

Finally, all surviving annotations for the start of unbridged repeats are marked as in-hinges and the annotations for the end of unbridged repeats are marked as out-hinges. One can formally show that under the assumption that no significant alignment is missed in the initial Overlap step, the contagion algorithm will place exactly one in-hinge and one out-hinge among the reads that originated from the set of unbridged occurrences of a repeat, and no hinge on the reads from the bridged occurrences of a repeat.

## HINGE-AIDED GREEDY ASSEMBLY ALGORITHM

A key distinction of HINGE’S approach to assembly lies in its graph layout step. Many OLC assemblers adopt the string graph paradigm (Myers 1995, 2005), which often produces assembly graphs that are unnecessarily dense. HINGE replaces the string graph algorithm with a variant of the greedy algorithm. This follows a recent line of work that found that variants of the greedy algorithm (such as the best-overlap-graph (BOG) algorithm (Miller et al. 2008b), “not-so-greedy” algorithm (Shomorony et al. 2016b) and the greedy merging algorithm (Shomorony et al. 2016a)) can produce a sparse overlap graph without mis-assemblies.

Notice that at the end of the contagion algorithm, we only have one in-hinge and one out-hinge for each unbridged repeat. In the graph layout step, we employ a variation of the greedy algorithm that utilizes the hinge information. Each read picks its left extension to be its longest prefix match, and its right extension to be the longest suffix match. However, unlike in the classical greedy algorithm, we do not restrict our search to overlaps. In addition to (prefix-suffix) overlaps, we also consider internal matches. Hence a read is allowed to find its successor/predecessor match to be an internal segment of another read, as long as the match starts on a hinge. An illustration of how internal matches are helpful in producing the correct graph layout, and a comparison with the classical greedy algorithm is illustrated in Supplemental Figure S2.

## THE ROLE OF POISONED READS

Another important aspect of the hinge-aided layout step is how the read poisoning information is used. As mentioned above, in the Contagion algorithm, whenever a read has its start/end repeat annotation killed by another overlapping read, we label it as poisoned. During the hinge-aided greedy algorithm, reads are prevented from picking a poisoned read as its predecessor/successor, guaranteeing that the two copies (or occurrences) of a bridged repeat remain separate.

This process is illustrated in Figure 6. In the scenario shown in Figure 6(a), read *u*_1_ is initially given a start-repeat annotation. However, this start-repeat annotation is removed by the contagion algorithm, as the repeat is bridged by read w. In this case, we keep a poisoned annotation on read *u*_1_. When a read has a poisoned start-repeat annotation, it cannot be chosen as a predecessor of another read if the match starts after the start-repeat annotation. As shown in Figure 6(b), according to a non-hinge-aided greedy assembly algorithm, *v*_2_ would choose *u*_1_ as its predecessor. However, as the match on *u*_1_ starts after the poisoned start-repeat annotation, we do not allow *v*_2_ to choose *u*_1_ as a predecessor. Instead we look for the next best option, which in this case is *u*_2_. This prevents a mis-assembly. The poisoning of end-repeat annotations works in an analogous way.

**Figure 6:**
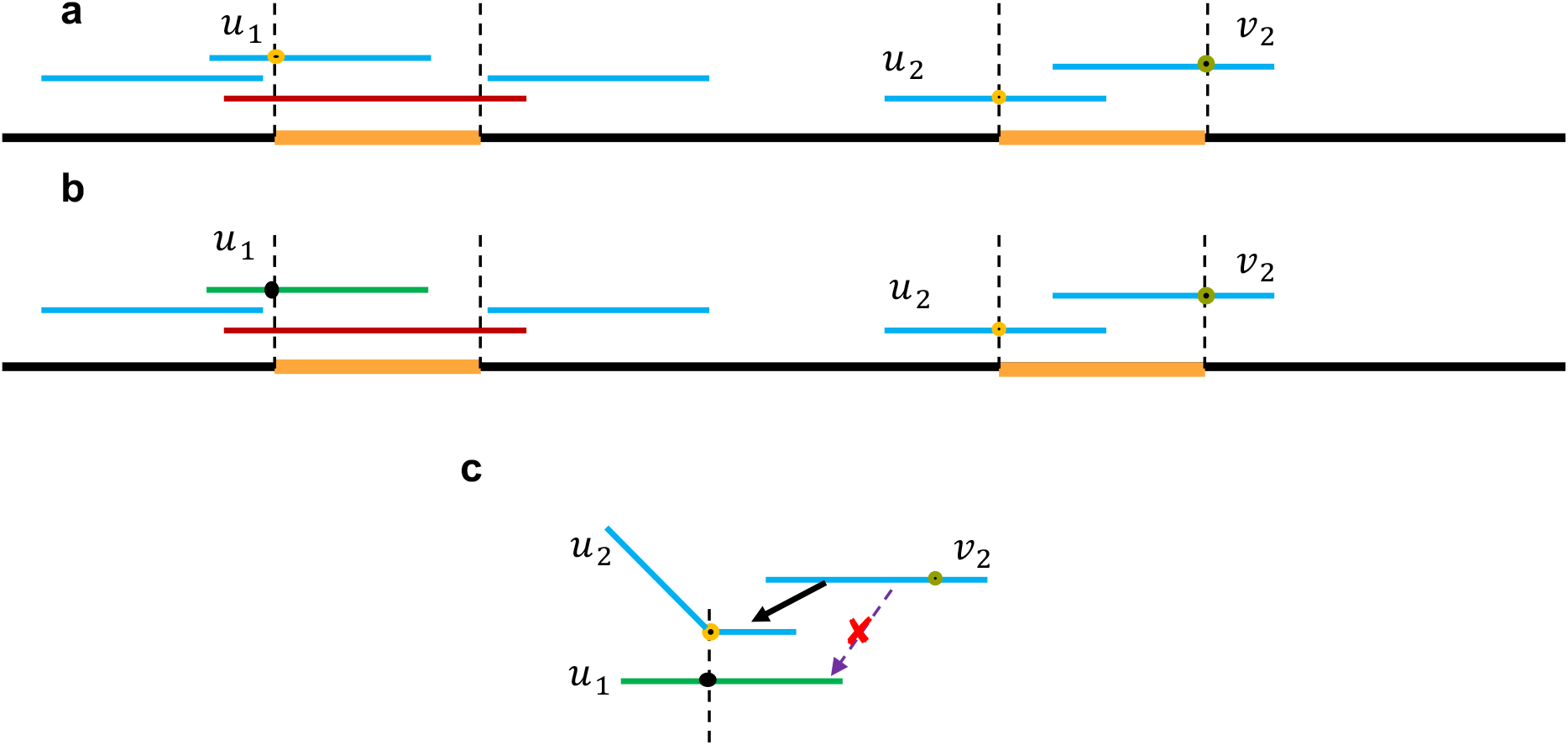
The *Poisoning* Algorithm: Read poisoning is part of the process by which we prevent a bridged repeat from collapsing on the graph, **(a)** In this scenario, read *u*_1_ is initially given a start-repeat annotation, which is killed in the contagion algorithm, as the repeat is bridged by read *w*. In this case, we keep a poisoned annotation on read *u*_1_. **(b)** When *v*_2_ looks for its best predecessor, it skips *u*_1_ due to the poisoned repeat annotation, preventing a mis-assembly.

In addition, we point out that the concept of poisoning is what allows the proper collapsing of the unbridged copies of repeats with three or more copies. Notice that during the third stage of the Contagion algorithm, we only consider non-poisoned reads. Therefore, we only deal with reads coming from unbridged copies of the repeat. As a result, the set of all unbridged occurrences of a repeats induces exactly one in-hinge and one out-hinge. On the other hand, all reads from the bridged copies are poisoned, and receive no hinge.

## REPEAT RESOLUTION

Another new ingredient introduced by HINGE is the use of global information to resolve repeats. Once constructed, the graph allows us to identify certain repeats that, although unbridged, can still be resolved based on the graph layout. As illustrated in Figure 3(a) and Supplemental Figure S15(a), when a repeat loop allows only one possible traversal, the loop can be untangled. We point out that the sparse and Eulerian-like nature of the graph produced by the hinge-aided greedy algorithm are important to allow this repeat resolution to be done in an automated fashion. We also point out that the loop resolution step is based on a parsimony principle, but it could be potentially incorrect if the loop corresponds to a plasmid, to a separate chromosome, or to the genome of a different species present in the sample. The parameter MAX_PLASMID_LENGTH sets the maximum length of a loop that should be considered a potential plasmid. HINGE will only resolve loops longer than MAX_PLASMID_LENGTH, and this behavior can be optionally turned off by setting MAX_PLASMID_LENGTH to a number longer than the genome length.

## HANDLING READ ORIENTATION AND DOUBLE-STRANDEDNESS

Since the orientation of the reads is unknown, as it is typical in all assembly pipelines one must consider each read and its reverse complement. Hence for each read we in fact create two nodes in the graph, and the constructed graph is symmetric. At the end of the graph construction, for visualization purposes, we overlay each node and its reverse complement.

## CONSENSUS

In order to generate consensus sequences for the resulting graph contigs, we first create a draft assembly by simply concatenating sections of the error-prone reads corresponding to unbranched paths on the graph. We then consider the alignment of all the original reads onto these draft contigs, and utilize a simple majority-based consensus to clean up these draft sequences. We reuse some code from FALCON (Chin et al. 2016) to perform this task. The result is output as a GFA file. We point out that the final contig sequences can be optionally run through Quiver (Chin et al. 2013) to further polish the assembly.

## GRAPH VISUALIZATION

All assembly graphs produced by HINGE were visualized using Gephi (Bastian et al. 2009).

## SOFTWARE AVAILABILITY

The HINGE assembler is available online at https://github.com/fxia22/HINGE and in the Supplemental Source Code. The analyses presented in Figure 2 can be reproduced in https://github.com/govinda-kamath/HINGE-analyses.

## ACKNOWLEDGEMENTS

The authors would like to thank Shoudan Liang and Jason Chin of Pacific Biosciences for useful discussions, and Lior Pachter of UC Berkeley for helpful comments and suggestions during the preparation of this manuscript. The authors are grateful to Nick Grayson and Julian Parkhill of The Wellcome Trust Sanger Institute for feedback and help with interpreting the results on the NCTC datasets. GMK would also like to thank John Lamping of Human Longevity Inc., chatting with whom drove him to take a data-driven approach to this project. Finally, GMK, IS, and FX would like to thank Gene Myers for presentations and work that were an inspiration to them, and for several tools that made this work possible.

## AUTHOR CONTRIBUTIONS

GMK and IS designed the algorithm. FX implemented a testbed to test and experiment with assembly algorithms. GMK, IS, and FX implemented the HINGE algorithm, ran it on the dataset, visualized and interpreted the results. TAC and DNT supervised the project. All authors wrote the paper.

